# RVPSD: A comprehensive and user-friendly web database for RNA viral protein structures

**DOI:** 10.64898/2026.02.06.704141

**Authors:** Qiangzhen Yang, Zhongshuai Tian, Tao Hu, Jiangrong Lou, Hengcong Liu, Edward C. Holmes, Yongyong Shi, Juan Li, Weifeng Shi

**Affiliations:** Bio-X Institutes, Key Laboratory for the Genetics of Developmental and Neuropsychiatric Disorders, Ministry of Education, Shanghai Jiao Tong University, Shanghai, China; Shanghai Institute of Virology, Shanghai Jiao Tong University School of Medicine, Shanghai, China; Department of Central Lab, The Second Affiliated Hospital of Shandong First Medical University, Tai’an, China; Key Laboratory of Emerging Infectious Diseases in Universities of Shandong, Shandong First Medical University and Shandong Academy of Medical Sciences, Ji’nan, China; School of Medical Sciences, The University of Sydney, Sydney, NSW 2006, Australia; School of Clinical and Basic Medical Sciences, Shandong First Medical University & Shandong Academy of Medical Sciences, Ji’nan, China; Ruijin Hospital, Shanghai Jiao Tong University School of Medicine, Shanghai, China; School of Life Sciences and Biotechnology, Shanghai Jiao Tong University, Shanghai, China

**Keywords:** RNA virus, protein structure prediction, Alphafold2, web database

## Abstract

Protein structures are central to understanding the diversity and biological function of RNA viruses. However, few experimentally validated structures of RNA viruses are currently available. To address this gap, we established the RNA Viral Protein Structure Database (RVPSD) that comprises diverse RNA viral protein structures predicted using AlphaFold2. RVPSD integrates taxonomic classification, viral nucleotide and protein sequences, functional annotations, and predicted three-dimensional protein structures to support a systematic exploration of the RNA viral proteomes. Currently, RVPSD comprises a total of 154,716 AlphaFold2-predicted RNA viral protein structures from 5,263 RNA virus species, with 84.9% exhibiting high pLDDT scores (>70) and representing a large-scale and comprehensive resource for RNA viral protein structures. RVPSD also provides a user-friendly web interface, with advanced search, sequence- and structure-based retrieval, interactive 3D visualization, and access and downloading structural data, enabling researchers to perform comparative structural analyses and functional annotation of viral “dark matter”.

## Introduction

RNA viruses are the most genetically diverse and rapidly evolving groups of microorganisms, causing a wide range of infectious diseases and posing severe threats to global health and biosecurity [1, 2]. Their high rates of mutation [3], sometimes frequent recombination [4], and highly compact genome organization [5] enable RNA viruses to rapidly evolve their protein functions through subtle genetic changes, facilitating adaptation to new hosts and environments, and often leading to the emergence or re-emergence of epidemic and pandemic variants [6]. In addition, a subset of RNA viral proteins, such as surface proteins and essential replication enzymes, have been identified as major targets for antiviral and vaccine development [7]. Understanding the three-dimensional (3D) conformations of RNA viral protein structures is therefore critical for elucidating the molecular mechanisms underlying viral infection, host adaptation, and immune evasion, as well as for guiding the rational design of antiviral therapeutics and vaccines [6, 8-11].

Experimental approaches such as X-ray crystallography and cryo-electron microscopy (cryo-EM) remain the gold-standard methods for resolving RNA viral protein structures [12]. However, achieving high-resolution structures via experimental methods is often hindered by the inherent physical properties of viral proteins, high conformational flexibility, and stringent biosafety requirements [12], leaving the number of experimentally determined RNA viral protein structures remarkably limited [13]. Consequently, computational structure prediction has emerged not only as a vital complement, but as a scalable alternative to experimental determination [14]. Recent breakthroughs in artificial intelligence (AI)–driven structure prediction, particularly AlphaFold, have revolutionized structural biology by enabling the high-throughput, atomic-level modeling of protein 3D conformations [15]. These advances have facilitated the generation of extensive data sets of protein models, such as the AlphaFold Protein Structure Database (AFDB) [16], that provide opportunities for systematic structural and functional analyses. Nevertheless, existing structural data are often dispersed across public repositories or restricted to individual virus families or specific virus proteins [17-19], thereby lacking a unified framework to facilitate comparative and evolutionary exploration across the entire RNA virosphere.

To bridge this gap, we developed the RNA Viral Protein Structure Database (RVPSD, https://virus.9itsg.net/#/home). This user-friendly web resource integrates over 150,000 highly accurate protein structures predicted using the AlphaFold2 pipeline across thousands RNA virus species classified by the International Committee on Taxonomy of Viruses (ICTV, https://ictv.global/). By providing a unified framework that connects structure, taxonomy, and sequences, RVPSD facilitates the systematic comparative analyses of viral evolution and virus-host interactions, which will in turn accelerate the rational design of antiviral agents.

## Results and discussion

### Overview of RVPSD

We developed RVPSD (https://virus.9itsg.net/#/home), an open-access platform integrating large-scale predicted protein structures across RNA virus taxa. RVPSD currently contains 154,716 predicted protein structures. All sequences are curated from the NCBI NCBI/GenBank database, followed by deduplication, taxonomic verification, and quality control. Each record in RVPSD encompasses viral species, strain, host range, genomic segment, and protein symbol, accompanied by AlphaFold2-predicted 3D models and associated metadata. Collectively, RVPSD establishes extensive resources currently available for structural and functional analysis of RNA viral proteins.

### Collection of RNA virus protein sequences

Using the dual-workflow strategy outlined in **Figure 1**, we assembled a data set comprising 154,716 RNA virus protein sequences. The ICTV-based workflow yielded a high-confidence protein data set comprising 20,448 protein sequences, derived from viral families formally curated in the ICTV VMR [20]. In parallel, the GenBank-based workflow contributed an additional 134,268 protein sequences from RNA viruses (realm *Riboviria*) that have not yet been included in the ICTV VMR.

**Figure 1.**
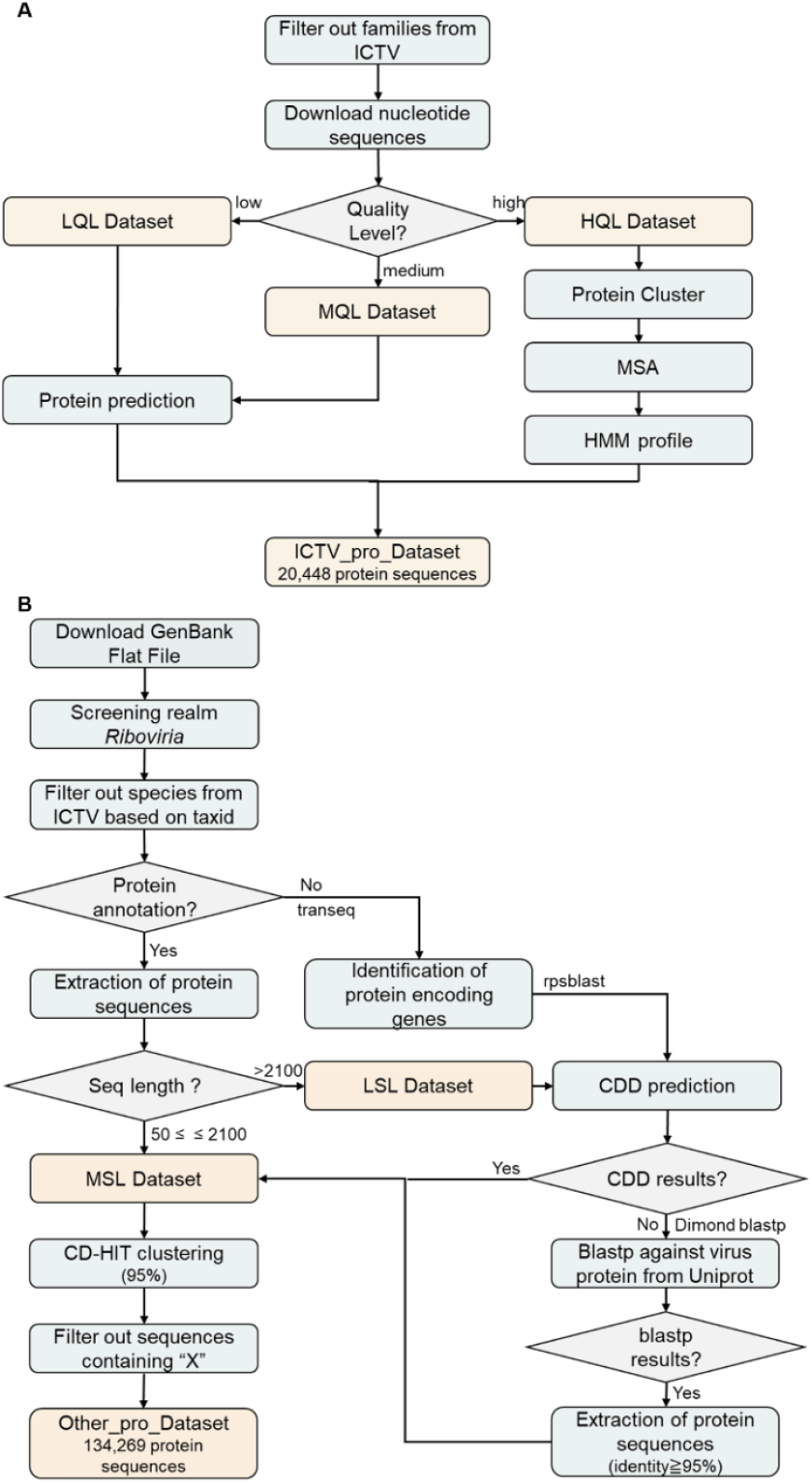
Overview of the workflows for viral protein sequence collection. (A) Viral protein sequences were collected through an ICTV-based workflow in which nucleotide sequences from ICTV-rectified viral families were processed, translated, and integrated into a unified protein data set. Based on annotation consistency, viral nucleotide sequences were stratified into high-, medium-, and low-quality categories (HQL, MQL, and LQL) data sets for downstream processing. (B) Viral protein sequences were independently collected through a GenBank-based workflow, in which sequences within the realm *Riboviria* were retrieved and curated through protein-coding gene identification, functional validation, and quality control procedures, including length filtering and redundancy removal. These workflows generated a final complementary protein data set that expands sequence coverage beyond ICTV-curated references.

Importantly, both workflows were anchored to the ICTV taxonomic system to ensure classification consistency. The ICTV-based workflow (**Figure 1A**) provides a taxonomically validated reference set, whereas the GenBank-based workflow (**Figure 1B**) extends sequence coverage to RNA viruses that have assigned taxonomy identifiers but have not yet been incorporated into the ICTV VMR. This design enables RVPSD to retain a stable taxonomic backbone while incorporating a broader spectrum of publicly available viral proteins, including those from emerging or less-characterized viruses [21]. Together, the final data set established a unified and systematically curated protein resource that ensures taxonomic reliability while maintaining sequence diversity.

### Taxonomic and Functional Coverage

Comprehensive analysis of the RVPSD data set demonstrates wide taxonomic and functional coverage. The database spans 93,211 nucleotide sequences of RNA viruses (based on nucleotide accessions) from 5,263 species and 149 families **(Figure 2A)**. Excluding *Retroviridae*, the families contributing the highest number of predicted protein models are the *Flaviviridae* (6.13%, 9,485/154,716), *Picornaviridae* (6.06%, 9,369/154,716), *Coronaviridae* (4.03%, 6,230/154,716), *Orthomyxoviridae* (3.72%, 5,758/154,716), and *Arteriviridae* (3.68%, 5,689/154,716), reflecting their extensive genomic resources **(Figure 2B)**. This distribution also reflects the prioritized sequencing efforts and extensive genomic resources dedicated to these families due to their impact on global public health and veterinary medicine [22, 23].

**Figure 2.**
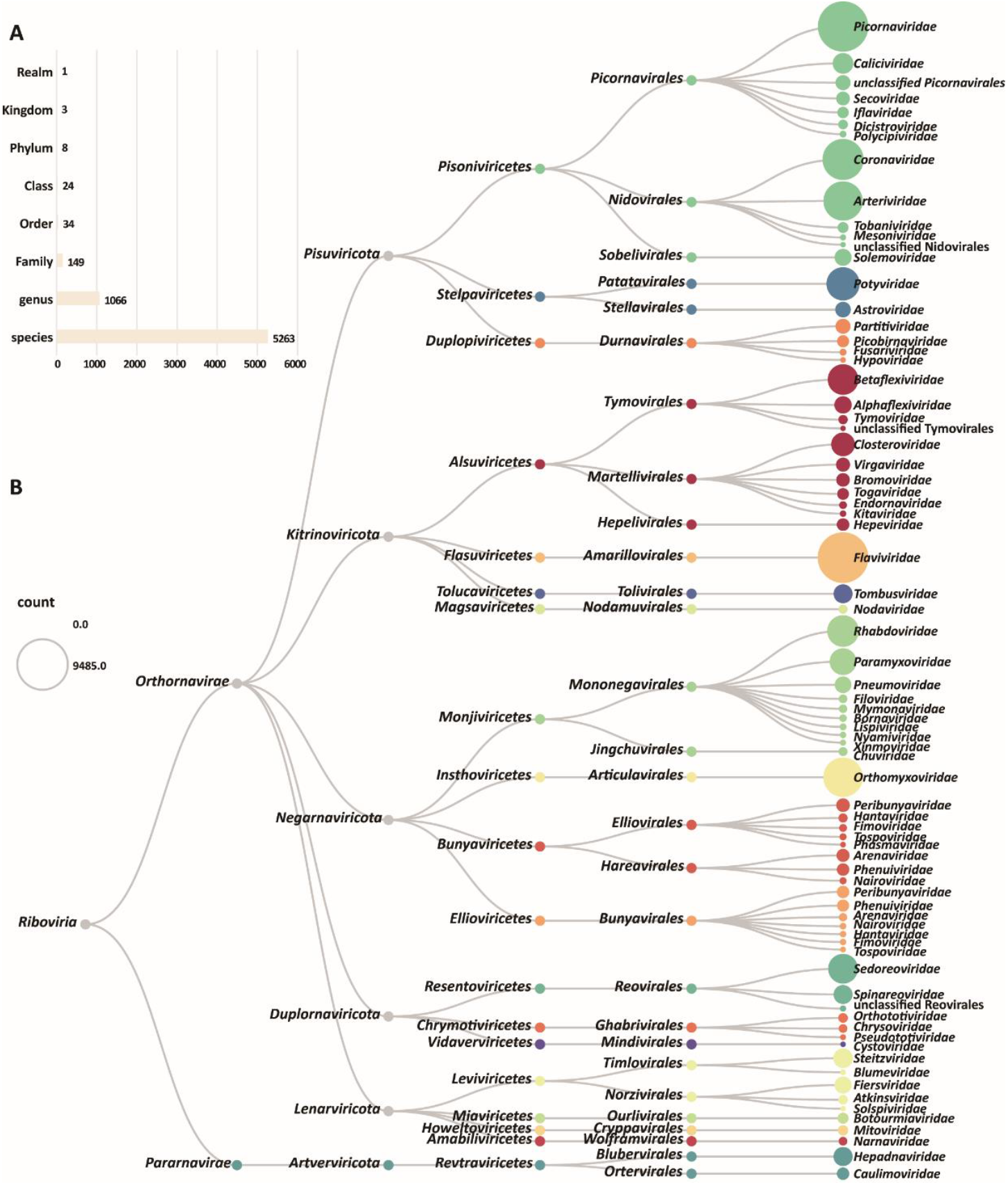
Taxonomic composition and hierarchical distribution of RNA viruses represented in RVPSD. (A)Numbers of taxonomic units represented in the RVPSD data set across major ICTV taxonomic ranks, comprising realm, kingdom, phylum, class, order, family, genus, and species. (B)The hierarchical taxonomic structure of RNA viruses included in RVPSD, shown from the realm *Riboviria* through the phylum, class, order, and family levels according to the ICTV classification framework. Only viral families represented by ≥100 RNA virus entries (based on nucleotide accessions) were shown, with *Retroviridae* excluded due to its distinct reverse-transcribing replication mechanism. Node size reflects the number of RNA virus entries within each lineage, and colors indicate different viral families to facilitate visual comparison of lineage-specific expansion patterns.

Functional annotation analysis was conducted on the subset of the proteins with assigned functional categories (n=139,599), excluding hypothetical or uncharacterized entries. Structural proteins (such as the capsid, envelope, and spike glycoproteins) constitute approximately 53.1% of the annotated entries, whereas nonstructural proteins (including polymerases, proteases, and helicases) account for 46.9%. This quantitative balance indicates that our data set contains sufficient protein structure data for subsequent use in the design of neutralizing antibodies and vaccines targeting structural proteins [24], as well as for the development of broad-spectrum antivirals targeting non-structural proteins such as the RNA-dependent RNA polymerase (RdRp) [25]. In addition, among proteins with host information (n=103,664), animal viruses represent 57.2% of entries followed by plant viruses (28.5%) and bacteriophage (14.3%). The inclusion of diverse host-associated viruses is essential for identifying conserved structural motifs that govern viral entry and replication across different biological kingdoms [26], thereby facilitating the study of the fundamental processes of RNA virus evolution and potential zoonotic spillover events.

### Predictive Modeling and Quality Assessment

The protein structures integrated into the RVPSD were predicted using the AlphaFold2 pipeline (**Figure 3A**). Model quality was evaluated using the predicted Local Distance Difference Test (pLDDT) score, a per-residue confidence metric that reflects the local accuracy of the predicted structures [15]. The cumulative pLDDT distribution across all predicted RNA viral proteins **(Figure 3B)** indicates the overall high model confidence. Specifically, 84.9% of the predicted structures achieved mean pLDDT scores greater than 70, 55.1% exceeded 80, and 21.7% exceeded 90, indicating reliable backbone prediction and high local structural accuracy. Notably, most structures showed pLDDT scores exceeding 70, which is commonly regarded as a threshold for high-confidence and accurate structural predictions [27].

**Figure 3.**
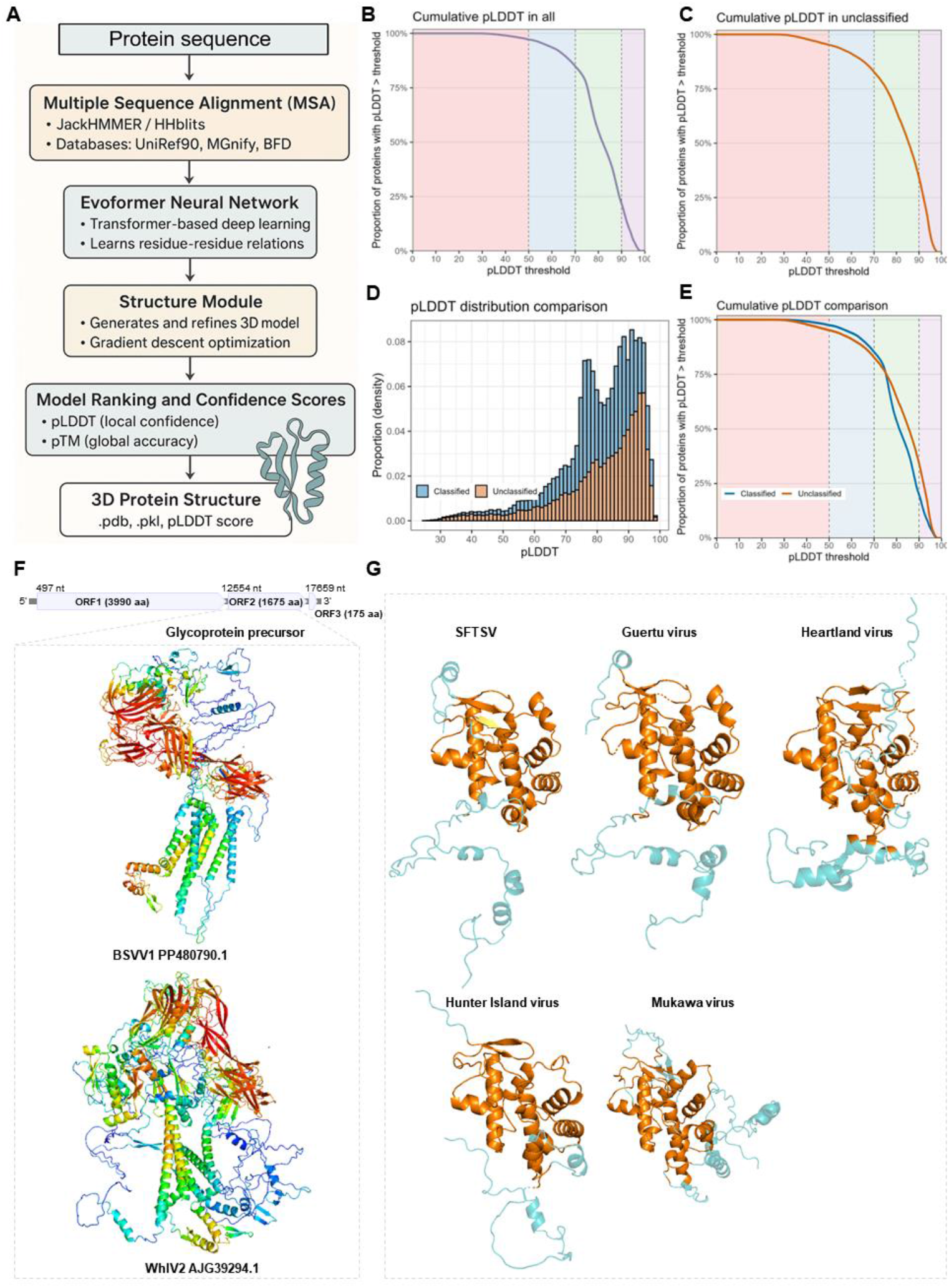
AlphaFold2-based structural prediction pipeline, pLDDT distributions, and web-based visualization in RVPSD. (A) Schematic overview of the AlphaFold2-based structure prediction workflow. (B) Cumulative pLDDT distribution for all predicted RNA viral proteins in RVPSD. (C) Cumulative pLDDT distribution for proteins from unclassified RNA viral taxa. (D) Comparison of the pLDDT distributions between proteins from classified and unclassified RNA viruses. (E) Cumulative pLDDT comparison between classified and unclassified protein sets. The curve represents the proportion of proteins with pLDDT greater than each threshold, illustrating differences in predicted structural confidence between these two groups. Shaded regions in (B), (C), and (E) indicate approximate interpretation ranges of pLDDT values: <50 (low confidence/likely disordered, red), 50–70 (moderate confidence, blue), 70–90 (reliable local structure, green), and >90 (high-confidence model, purple). (F) Representative sequence-based functional inference using RVPSD. AlphaFold-predicted structure of BSVV1 ORF2 [31] (top), and representative viral glycoprotein structures retrieved from RVPSD based on sequence similarity (bottom), both colored by per-residue pLDDT scores. (G) Structure-based comparison of NSs from multiple bunyaviruses. NSs protein structures corresponding to sequences with detectable amino acid similarity to SFTSV NSs were retrieved and downloaded from RVPSD. Accession number of SFTSV NSs, Guertu virus NSs, Heartland virus NSs, Hunter Island virus NSs, and Mukawa virus NSs were AVV65325.1, QBQ64951.1, UDE22460.1, AHI10996.1, and BBE15817.1, respectively. Structurally overlapping regions identified from the multi-structure comparison are highlighted (orange).

To assess whether prediction quality varied with taxonomic annotation status, pLDDT distributions were also examined for proteins from unclassified RNA viral taxa **(Figure 3C)** and directly compared with those from taxonomically classified viruses **(Figures 3D and 3E)**. The cumulative and density-based comparisons revealed highly consistent confidence profiles between the two groups. Specifically, 85.2% of the classified proteins and 82.8% of the unclassified proteins achieved pLDDT scores >70, confirming that the AlphaFold2 prediction accuracy is robust and independent of the classification status of the virus species. The consistency also validates that our strategy successfully captures high-quality structural data even for emerging or previously uncharacterized viral lineages. Consequently, these results validate the overall high quality of the RVPSD data set, supporting its utility for large-scale structural and functional analyses of RNA viruses, including potentially “novel” species.

### Database Architecture and Functional Implementation

To enable efficient access to and exploration of the large-scale RNA viral protein structure data set generated in this study, we implemented RVPSD as a web-based platform with an integrated backend system (**Supplementary Methods**). Building on this infrastructure, RVPSD provides a unified and user-oriented interface for sequence- and structure-based retrieval, visualization, and output management of RNA virus protein structures (**Figure S1**).

The platform is organized into five functional modules: (i) Home, (ii) Advanced Search, (iii) Sequence Search, (iv) Structure Search, and (v) Query History. The Home module presents overall database statistics, update information, and external resource links. The Advanced Search module supports flexible querying using accession numbers, species names, protein symbols, or host taxonomy, with search results summarized in tabular form **(Figure S1A)**. Sequence-based retrieval is implemented through the integration of BLASTP [28], allowing users to submit protein sequences and identify the most similar entries in RVPSD. Search results are displayed with detailed alignment statistics, including alignment length, start position, and Bit score **(Figure S1B)**, and can be accessed either through the Query History module or via automated email delivery. Structure-based retrieval is enabled through Foldseek [29], which allows users to upload protein structure files and identify structurally similar models within RVPSD. Results are ranked by TM-score and presented alongside structural thumbnails and associated transformation parameters.

For all query modes, individual search results link to a dedicated detail page that provides comprehensive contextual information, including GenBank accession numbers, taxonomic classification, strain information, and host annotation. Predicted 3D protein structures are visualized interactively using the Mol* viewer [30], enabling residue-level inspection and the direct download of the corresponding PDB files (**Figure S1C**). Collectively, these functionalities allow RVPSD to support efficient data retrieval, interactive structural inspection, and exploratory analysis of RNA viral protein structures through a single, integrated web interface.

### Examples of the utility of RVPSD

#### Case I: Functional inference of a “dark” viral protein using RVPSD

Brine shrimp virga-like virus 1 (BSVV1) is a recently identified RNA virus with an unusual genome organization potentially derived from an ancient recombination between the positive-sense (*Martellivirales*) and the negative-sense (*Bunyavirales*) RNA viruses [31]. Its ORF2 encodes a highly divergent protein with limited amino acid sequence identity (31.92%) to known viral proteins, complicating functional inference based on conventional sequence analysis alone. To explore its potential function, the amino acid sequence of BSVV1 ORF2 was first subjected to *de novo* structure prediction using AlphaFold3 (https://alphafoldserver.com/), which yielded a folded model with a mean pLDDT score of 57.0 (**Figure 3F**). The ORF2 sequence was then queried against the RVPSD database using the sequence retrieval module. RVPSD identified viral glycoproteins from Wǔhàn insect virus 2 (WhIV-2, *Bunyavirales*) as the top-ranked hit, despite a low sequence identity of 29.2% (**Figure 3F**). Structural comparison indicated broadly similar overall structural organization, with comparable arrangements of major α-helical regions between BSVV1-ORF2 and known bunyavirus glycoproteins (structure-based multiple structural alignment LDDT = 0.516). These observations were consistent with previous evidence [31], confirming that BSVV1 ORF2 is evolutionarily related to the glycoproteins of negative-sense RNA viruses. This case illustrates how RVPSD can facilitate functional inference for highly divergent viral proteins.

#### Case II. Structure-based identification of conserved regions in bunyavirus NSs

The non-structural protein (NSs) of severe fever with thrombocytopenia syndrome virus (SFTSV) is a key virulence factor involved in interferon signaling, inflammatory responses, RNA interference, and autophagy pathways [32]. However, pronounced sequence divergence among bunyavirus NSs proteins hampers sequence-based identification of shared features, making structure-resolved resources such as RVPSD essential for comparative analysis. NSs structures corresponding to sequence-similar hits of SFTSV NSs were retrieved and downloaded from RVPSD, including representatives from Guertu virus, Heartland virus, Hunter Island virus, and Mukawa virus, which demonstrated 21.6–75.5% amino acid sequence identities to the SFTSV NSs. Multi-structure comparison of the retrieved NSs proteins revealed the overlapping regions shared by the five viruses, primarily corresponding to the compact α-helical cores **(Figure 3G)**. Consistent with this observation, multiple structural alignment yielded an average multiple structural alignment LDDT score of 0.610, indicating a moderate structural consistency despite substantial sequence divergence. Several motifs in SFTSV NSs have been previously reported to play a critical role in viral immune evasion and pathogenesis, including two conserved sequence motifs (SALRWPSG and DWP) and two predicted protein– protein interaction motifs (PKNP for SH3-domain interaction and WRGL for LC3 interaction) [33]. Apart from the DWP motif, the remaining three motifs were spatially located within the overlapping structural core we identified (**Figure S2**), suggesting that many other bunyaviruses may also employ these function motifs in the NSs protein for immune evasion. This example illustrates how RVPSD enables downstream structure-based comparative analyses, facilitating the identification of shared structural features that are not readily apparent from sequence comparison.

Overall, RVPSD represents a comprehensive and user-friendly web database that integrates predicted 3D structures of RNA viral proteins across diverse viral lineages. By combining taxonomic classification with curated sequence information and high-confidence AlphaFold2 structural models, RVPSD provides a reliable platform for structural prediction, inspection, retrieval, and visualization of RNA virus proteins. While the current implementation focuses on sequence- and structure-based search and model exploration, the breadth and quality of the data set contribute to enabling future detailed structural comparisons across RNA virus proteins. As viral sequence resources continue to expand and structure prediction methodologies advance, RVPSD will be regularly maintained and updated.

## Conclusion

By integrating large-scale protein sequences, AI-driven predicted structural data, comprehensive functional annotations and analytical tools, RVPSD provides a unified and user-friendly platform for a wide range of virological applications, and may contribute to the development of effective countermeasures against emerging viral threats.

## Methods

RVPSD was developed to support large-scale structural analysis of RNA viruses by integrating viral protein sequences and predicted structures. We employed a dual-workflow strategy to assemble the data set: an ICTV-based workflow and a GenBank-based workflow. Viral nucleotide sequences were collected according to the ICTV taxonomy [20] or retrieved from the National Center for Biotechnology Information (NCBI) GenBank FTP server [21] and curated into standardized protein data sets. Protein structures were predicted using AlphaFold2 [15], and confidence metrics were extracted for downstream analysis. The RVPSD platform was implemented with a Vue3based web interface and a MySQL backed database, enabling efficient data access, visualization, and sequence or structure-based analysis. For representative use cases (Case I and Case II), protein structures were retrieved from RVPSD, and subjected to structure-based multiple alignment using FoldMason [34], followed by visualization in PyMOL. Detailed data processing pipelines, quality control procedures, and workflow design are provided in the supplementary material.

## Author contributions

Weifeng Shi and Yongyong Shi conceived and supervised the project. Juan Li, Zhongshuai Tian, and Tao Hu were responsible for constructing the dataset. Qiangzhen Yang, Hengcong Liu, and Jiangrong Lou were responsible for predicting RNA viral structures. Juan Li, Qiangzhen Yang, Zhongshuai Tian, and Tao Hu analyzed the results, wrote the manuscript, and generated the figures. Weifeng Shi, Juan Li, Yongyong Shi, and Edward C. Holmes revised the manuscript. All authors read and approved the final manuscript.

## Acknowledgement

We thank the researchers who generated and shared the sequencing data from the NCBI GenBank database. The computations in this paper were run on the Siyuan-1 cluster supported by the Center for High Performance Computing at Shanghai Jiao Tong University.

This work was supported by the Fundamental Research Funds for the Central Universities (YG2023ZD01), the National Natural Science Foundation of China for Distinguished Young Scholars (32325003 to W.S.), the National Natural Science Foundation of China (32370724 and 82472254), the National Key R&D Program of China (2023YFC2307503), the Science and Technology Commission of Shanghai Municipality (2023ZR1436300 and 23JC1403400), and the Natural Science Foundation of Shandong Province (ZR2024MH288). J.L. is supported by the Taishan Scholars Programme of Shandong Province (tsqn202211217) and the National high-level talents special support plan - Ten thousand plan - Young top - notch talent support program. E.C.H. is supported by a National Health and Medical Research Council (NHMRC, Australia) Investigator Award (GNT2017197).

## Data and code availability

All viral sequence data and associated metadata used in this study were sourced from publicly accessibly NCBI databases. The nucleotide and protein data set comprised all sequences from the realm *Riboviria* (NCBI Taxonomy ID: 2559587) deposited in NCBI up to October 23, 2023. The taxonomic framework for viral classification was based on the ICTV Virus Metadata Resource (VMR_21-221122_MSL37). All data sets and scripts have been deposited on the RVPSD website (https://virus.9itsg.net/#/home). Supplementary materials (methods, figures, graphical abstract, and updated materials) may be found in the online DOI or iMeta Science http://www.imeta.science/.

## Competing interests

The authors declare no competing interests.

## Ethics statement

No animals or humans were involved in this study.

---

**Figure S1.**
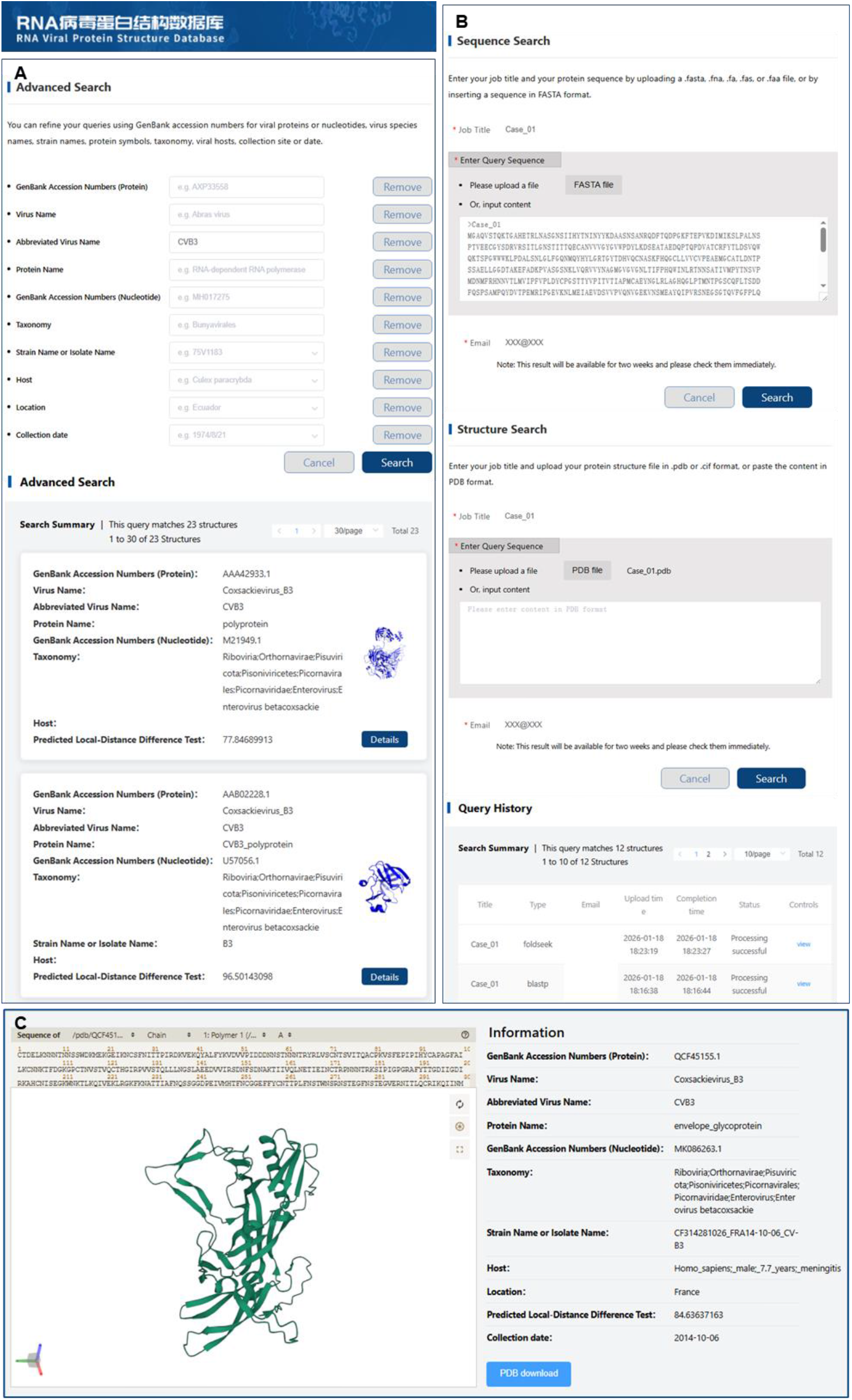
Web interface and search functions of RVPSD. (A)Advanced Search module, enabling refined queries using accession numbers, virus names, taxonomy, host information, and collection metadata, with results summarized in tabular format. (B) Search and retrieval interfaces, including sequence-based search via BLASTP, structure-based search via Foldseek, and the Query History module displaying submitted jobs and corresponding retrieval results. (C) The entry visualization page provides an interactive 3D structure viewer alongside detailed metadata of each hit, including sequence information, GenBank accession, taxonomy, and a direct link for structure download.

**Figure S2.**
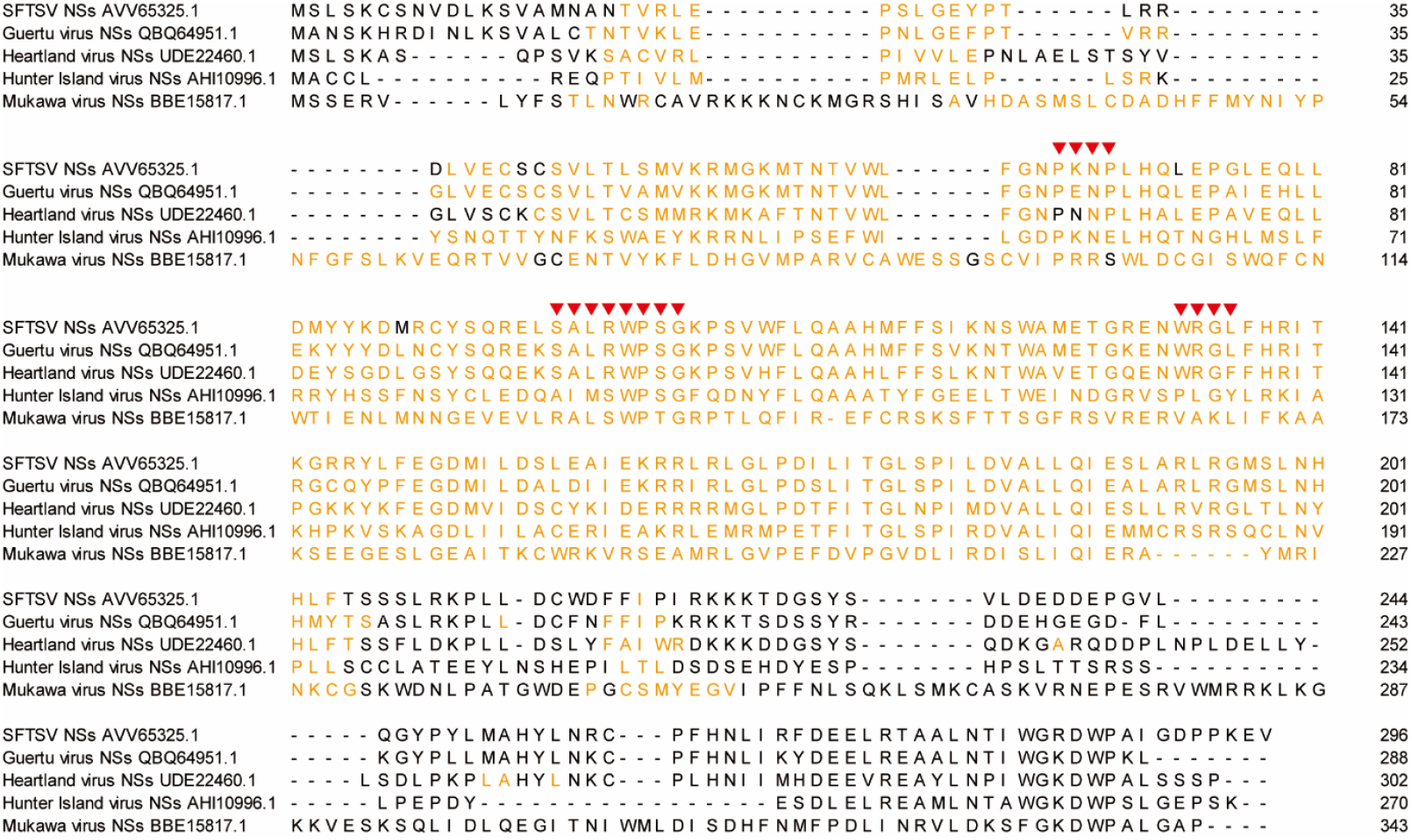
Sequence alignment of bunyavirus NSs. Multiple sequence alignment of NSs proteins from SFTSV, Guertu virus, Heartland virus, Hunter Island virus, and Mukawa virus corresponding to the structure-based comparison shown in Figure 3G. Residues highlighted in yellow indicate positions mapping to the shared structural overlap/core. Red triangles denote previously reported SFTSV NSs motifs located within this region.

## Supplemental information

## Methods

### ICTV-based protein sequence collection

Viral nucleotide sequences were collected at the family level according to the International Committee on Taxonomy of Viruses (ICTV, 2024 release) to establish a taxonomically standardized data set [1]. To ensure both quality and completeness, sequence quality assessment was conducted by integrating three independent information sources: the ICTV Virus Metadata Resource (VMR, VMR_21-221122_MSL37), the National Center for Biotechnology Information (NCBI) GenBank annotations [2, 3], and published literature.

For each viral nucleotide sequence listed in the ICTV VMR, annotation consistency was evaluated by comparing NCBI GenBank annotations against ICTV classifications and literature reports. Based on this comparison, sequences were assigned to three quality levels:(i)sequences with annotations that were consistent across NCBI annotations, ICTV classification, and literature reports were classified as “high-quality” level (HQL); (ii) sequences with annotations consistent between NCBI annotations and either the ICTV classification or literature reports were classified as “medium-quality” level (MQL); and (iii) sequences with NCBI annotations inconsistent with both the ICTV classification and literature reports were classified as “low-quality” level (LQL).

Nucleotide sequences classified as MQL and LQL lacked complete or reliable protein annotations and were therefore subjected to protein prediction to infer putative protein-coding regions. In contrast, HQL sequences exhibited well-defined coding regions and high assembly quality, enabling direct translation into protein sequences. Proteins derived from HQL sequences were further processed through protein-level clustering, multiple sequence alignment (MSA), and hidden Markov model (HMM) profile [4] construction to capture the conserved evolutionary features across viral protein families. Protein sequences obtained from all quality tiers were integrated into a preliminary ICTV-based protein prediction data set, and the hierarchical visualization of RNA virus taxonomy was generated using RAWGraphs [5].

### Supplementary protein sequence collection from non-VMR viral species

To complement the ICTV-curated protein sequence data set, additional viral protein sequences derived from viral species not included in the ICTV VMR were collected through an independent GenBank-based analysis workflow. All viral nucleotide sequences were downloaded from the NCBI GenBank FTP server (https://ftp.ncbi.nlm.nih.gov/genbank/, Release 249.0) [2], and entries belonging to RNA viruses within the realm *Riboviria* were retained. Based on the taxonomic identifiers, viral species absent from the VMR file were identified using the ETE3 NCBITaxa python package [6] and subsequently classified according to the presence or absence of protein annotations.

For viral sequences containing protein annotations, protein sequences were extracted using the Bio.SeqIO module (https://biopython.org/wiki/SeqIO) available in Biopython v1.43 onwards [7], and sequences shorter than 50 amino acids were discarded. Because protein sequences longer than 2,100 amino acids were not compatible with subsequent structure prediction, different processing strategies were applied based on sequence length. Protein sequences shorter than 2,100 amino acids were directly retained for downstream processing. Protein sequences longer than 2,100 amino acids were searched against the Conserved Domain Database (CDD) [8] using rpsblast from BLAST+ v2.6.0+ (E-value ≤1e−10) [9], and conserved regions were extracted to retain protein fragments with lengths between 50 and 2,100 amino acids.

For viral sequences lacking protein annotations, protein-coding regions were predicted using EMBOSS transeq [10]. Predicted protein sequences were first searched against the CDD using rpsblast (BLAST+ v2.6.0+, E-value ≤1e−10). Protein regions with detectable conserved domains were retained for downstream processing. For predicted protein sequences without detectable CDD annotations, additional similarity searches were performed using BLASTP v2.6.0+ [9] against the UniProt viral protein database (uniprot_trembl_viruses.dat.gz, 2023-09-13) [11], and sequences with identity ≥95% were retained.

All retained protein sequences were subjected to standardized post-processing, including length filtering (50-2,100 amino acids), removal of sequences containing ambiguous amino acid residues (“X”), and redundancy elimination using CD-HIT v4.7 (parameters: -c 0.95 -n 5 -M 16000) [12]. The resulting sequences were integrated into the GenBank-derived protein data set for downstream structural analyses.

### Protein structure prediction

Three-dimensional structures of the RNA viral proteins were predicted using AlphaFold v2.0 [13] with default parameters. Briefly, the amino acid sequence of each protein was used as input. MSAs were generated against major sequence databases (UniRef90 [14], Mgnify [15], and BFD (https://bfd.mmseqs.com/)) using JackHMMER [4] or Hhblits [16], capturing co-evolutionary relationships that inform residue proximity and structural constraints. The Evoformer module integrated MSA and template information to infer inter-residue distances and orientations, which were further refined by the structure module through iterative gradient-based optimization. Final models were ranked according to the predicted Local Distance Difference Test (pLDDT) score, reflecting per-residue confidence [13].

All predicted protein structures were processed using Biopython v1.83 [7] for residue indexing and standardized into PDB and mmCIF formats. Structural metadata of each protein entry, including model confidence, residue coverage, and secondary structure composition, were extracted using PyMOL v2.6 [17] and DSSP v4.3.0 [18] to support database integration.

### Database construction, system architecture, and integrated analytical tools

RVPSD was developed on a classical three-tier web–application–database architecture. The front-end interface was implemented using Vue3 (https://vuejs.org/) and JavaScript, ensuring responsive performance and intuitive visualization. Backend services were developed in Java using the Spring Boot framework, with an embedded Tomcat server (https://tomcat.apache.org/) responsible for data parsing, search handling, and result generation. All data were stored in a MySQL 8.0 (https://dev.mysql.com/) relational database and optimized through indexed tables to improve query efficiency. The application was deployed on a Linux server environment (AlmaLinux 8.10) (https://almalinux.org/blog/2024-05-28-announcing-810-stable/) and configured with an Nginx reverse proxy to ensure stable and reliable user access.

To support sequence- and structure-based exploration of the RNA viral proteins, the RVPSD platform integrates multiple analytical tools. Protein-level sequence similarity searches are supported using BLASTP [9], while structure-based retrieval is enabled through Foldseek [19] for efficient structural comparison. Query results are recorded in the Query History module and can be accessed by users for result tracking, with selected results automatically delivered to user-specified email addresses. Interactive three-dimensional visualization of viral protein structures was enabled through the Mol* viewer [20], supporting dynamic rendering of the AlphaFold2-predicted models and allowing users to rotate and inspect protein structures at the residue level directly within the browser.

### Structure-based comparative analysis and visualization of RVPSD-derived proteins

Protein structures used for structure-based comparative analyses were retrieved from the RVPSD web interface. For Case I and Case II, representative protein structures identified through sequence- or structure-based retrieval were downloaded and subjected to external structure comparison. Multi-structure alignment was performed using FoldMason [21] with default parameters. Structural visualization and identification of the overlapping regions were performed using PyMOL v2.3.2 (https://pymol.org/). These analyses were performed as downstream applications to illustrate the utility of RVPSD-derived structural data.

### Data availability

All predicted protein structures, annotations, and visualization data were freely available through the RVPSD website (https://virus.9itsg.net/#/geneContrast). The individual protein structures in PDB files can be downloaded under the “Downloads” module. Source codes for data processing and web deployment were available upon request for academic use.

## Notes

### Competing Interest Statement

The authors have declared no competing interest.

